# Transcriptional-Regulatory Convergence Across Functional MDD Risk Variants Identified by Massively Parallel Reporter Assays

**DOI:** 10.1101/2021.03.05.434177

**Authors:** Bernard Mulvey, Joseph D. Dougherty

**Affiliations:** Depts. of Genetics and Psychiatry, Washington University in St. Louis

**Author notes:** Corresponding Author: Joseph D. Dougherty, Depts of Genetics and Psychiatry, Washington University in St. Louis, Address: 660 S. Euclid Ave, Campus Box 8232, St. Louis, MO 63110.

**Keywords:** Depression, Massively Parallel Reporter Assay, Retinoid Signaling, Omnigenic, Functional Annotation

## Abstract

Family and population studies indicate clear heritability of major depressive disorder (MDD), though its underlying biology remains unclear. The majority of single-nucleotide polymorphism (SNP) linkage blocks associated with MDD by genome-wide association studies (GWASes) are believed to alter transcriptional regulators (e.g., enhancers, promoters), based on enrichment of marks correlated with these functions. A key to understanding MDD pathophysiology will be elucidation of which SNPs are functional and how such functional variants biologically converge to elicit the disease. Furthermore, retinoids can elicit MDD in patients, and promote depressive behaviors in rodent models, acting via a regulatory system of retinoid receptor transcription factors (TFs). We therefore sought to simultaneously identify functional genetic variants and assess retinoid pathway regulation of MDD risk loci. Using Massively Parallel Reporter Assays (MPRAs), we functionally screened over 1 000 SNPs prioritized from 39 neuropsychiatric trait/disease GWAS loci, with SNPs selected based on overlap with predicted regulatory features—including expression quantitative trait loci (eQTL) and histone marks—from human brains and cell cultures. We identified >100 SNPs with allelic effects on expression in a retinoid-responsive model system. Further, functional SNPs were enriched for binding sequences of retinoic acid-receptive transcription factors (TFs); with additional allelic differences unmasked by treatment with all-*trans* retinoic acid (ATRA). Finally, motifs overrepresented across functional SNPs corresponded to TFs highly specific to serotonergic neurons, suggesting an *in vivo* site of action. Our application of MPRAs to screen MDD-associated SNPs suggests a shared transcriptional regulatory program across loci, a subset of which are unmasked by retinoids.

## INTRODUCTION

Major depressive disorder (MDD) is a common but debilitating psychiatric disorder affecting hundreds of millions worldwide^1^, exacting substantial tolls on both individuals^2^ and societies^3^. Despite the global burden of MDD, nearly half of patients do not clinically respond to treatment^4^, in part due to limited understanding of its biological underpinnings. Family studies have demonstrated that MDD risk is at least 30% heritable^5,6^. More recently, genome-wide association studies (GWASes) have demonstrated similar estimates for severe MDD^7^, and have helped narrow in on associated single nucleotide polymorphisms (SNPs)^8–12^, a tantalizing entry point for understanding the biology of MDD. However, GWAS studies do not identify causal variants, but rather implicate wider co-inherited regions consisting of many SNPs in linkage disequilibrium (LD). Most disease-associated SNPs are found outside of protein-coding sequences suggesting probable roles in transcriptional regulation (TR)^13–16^. However, this leaves unresolved issues of identifying which linked SNPs have functional allelic impacts on TR, and how these act together across loci to result in disease.

In addition, it is generally thought that undetected, small-effect SNPs acting across the genome—including conditional SNPs within GWAS-significant loci^17^—contribute to the substantial heritability *not* caught by GWAS-significant SNPs alone^18^. A recent analysis of psychiatric disorder GWASes confirmed the existence of such conditional SNPs in MDD^19^. As such, multiple SNPs with allelic TR effects could exist per LD block, but functional demonstration has been sparse to date. The largest functional TR assay of MDD-associated variants examined 34 SNPs using luciferase assays^20^, representing successful but low-throughput identification of functional MDD SNPs. However, in terms of broad linkage, these loci constitute well over 10,000 SNPs, which will ultimately require higher-throughput approaches.

Furthermore, how functional SNPs biologically result in disease remains unclear, even once such SNPs are identified. Nonetheless, polygenic^21^ and omnigenic^18^ models provide a guiding principle for interpretation of complex disease genetics. In brief, these theories posit that consistent emergence of a specific phenotype via widespread genomic variation necessarily requires common biological endpoints of those variants’ effects. At the molecular level, these points of convergence could be either upstream (shared regulation across loci)^22^ or downstream (common biological pathways across loci). For downstream analyses, myriad approaches have been developed to nominate gene targets of putative TR SNPs using proximity^23^, chromatin structure^24–26^, or expression quantitative trait loci (eQTLs)^27–29^, yielding gene sets tested for enrichment in biological pathways^27,28,30^ and cell types^31^. However, no analogous approaches exist for identifying convergent upstream (*i.e.,* TR) molecular effects of genetic risk, in part because a prerequisite is defining the functional SNPs.

One possible point of upstream TR convergence of MDD risk variants is retinoic acid and related compounds (retinoids). Retinoids are steroid hormones, driving transcriptional responses via several retinoid-binding transcription factors (TFs) and heterodimerizing partners^32,33^. Besides their critical role in neurodevelopment, including of depression-implicated limbic structures^34^, retinoids have been associated with MDD onset and suicidality by epidemiological studies of the retinoid agonist isotretinoin^35^. Additional evidence for retinoid pathway activity in the adult brain—and its overactivity as a risk factor for depression—comes from rodent pharmacology and genetic models. For example, knockdown of *Cyp26b1*—which metabolizes retinoids—in adult mouse anterior insula suppresses interest in social novelty by reducing spontaneous activity of excitatory neurons^36^. Likewise, depressive symptoms have been observed in rats after intracerebroventricular all-*trans* retinoic acid (ATRA) administration^37^. In addition, retinoid agonism of the TF RARA upregulates corticotrophin releasing hormone (*CRH*) expression, and RARA is more abundant in CRH neurons of affective disorder hypothalami^38^; moreover, RARA prevents corticoid negative feedback on CRH expression^39^, resulting in overactivity of stress signaling in the brain and body. Finally, given the substantial shared genetic risk across psychiatric disorders^40^, it is notable that schizophrenia GWAS loci show enrichment for retinoid TR^41^,and that circulating retinoids are dysregulated in schizophrenia patients ^42^. Similarly, retinoid pathway genes including *CYP26B1* are dysregulated in postmortem brain from autism spectrum disorders, bipolar disorder, and schizophrenia patients^43^. These findings led us to speculate that a component of MDD risk genetics may, in fact, demonstrate an upstream convergence via recurrent retinoid-mediated TR disruptions by MDD-associated SNPs.

Massively parallel reporter assays (MPRAs) provide a solution to both experimentally identify functional variants and, consequently, their shared TR features. MPRAs assess thousands of DNA elements for transcriptional-regulatory functions and allelic differences simultaneously by pairing each short genomic sequence element of interest to several unique barcodes, with a constant promoter and reporter gene placed in between^44–47^. Delivery of a library of DNA elements to cells, followed by RNA collection and sequencing, enables quantitative estimation of the expression driven by each element as a ratio of collected RNA barcode to delivered DNA barcode. These assays have recently been adapted to systematically identify SNPs with functional allelic TR differences from GWAS loci for several diseases^48–56^. Two key features make MPRAs advantageous for identifying both functional SNPs and their TR interactions. ^57^First, the assay is carried out via transfection and targeted RNA-sequencing, meaning it can be executed in unmodified cell lines appropriate to the application. Second, MPRAs can be conducted to define TR effects of experimental manipulations in these systems, such as steroid hormone administration^57^.

Therefore, we sought to experimentally identify functional TR SNPs from 39 GWAS loci associated with MDD, neuroticism, and broader psychiatric disease risk, with the hypothesis that functional SNPs converge at the level of retinoid-mediated TR. From broad linkage regions, we selected over 1 000 SNPs based on overlapping human brain and neural epigenomic signals suggestive of TR activity. Critically, selection of neither the loci nor the SNPs was predicated on retinoid involvement, allowing for unbiased functional screening of a cross-section of MDD GWAS loci. To ensure we could detect SNPs subject to retinoid-mediated TR, we used neuroblastoma (N2a) cells, as they are strongly and rapidly retinoid-responsive^58,59^. Our initial assay identified over 75 functional SNPs from 29 GWAS regions, confirming that GWAS loci contain several functional SNPs. We then examined whether these functional SNPs possessed shared upstream TR features— namely, transcription factor (TF) binding motifs. Remarkably, there was indeed enrichment of retinoic acid binding TFs among the MPRA-functional vs. -nonfunctional SNPs, supporting our hypothesis. To further characterize retinoid effects on TR at MDD-associated SNPs, we performed a second assay using all-*trans* retinoic acid (ATRA), known to stall division of N2a and other neuroblastoma cells by inducing neuronal-like differentiation^58^. First, we found that functional SNPs containing retinoid receptor motifs had increased magnitudes of effect in the presence of ATRA, consistent with bonafide retinoid receptor TR activity. More broadly, ATRA led to striking rearrangements of the baseline regulatory landscape, including altered magnitude and reversed direction of allelic effects. Additionally, it revealed new SNPs with allelic TR differences unmasked by ATRA treatment. Significant ATRA-allele interaction SNPs were enriched not only for binding sequences of the retinoid receptor RARG, but for motifs of several known retinoid-induced TFs, indicating broad roles of both retinoid TFs and their downstream TR systems at functional MDD-associated SNPs.

Finally, we explored the cell type-specificity of TFs predicted to regulate our functionally identified SNPs. Strikingly, we found TFs highly specific to serotonin neurons were strongly enriched among those we predicted to be recurrently involved in retinoid-dependent SNP function. These findings suggest that the broad transcriptional-regulatory systems engaged by retinoids—and as we illustrate, the genetic component of MDD risk *they* engage—may converge on serotonergic neurons. In summary, we identify MDD-associated functional SNPs with both baseline and ATRA-mediated allelic differences in TR, and these disproportionately show upstream convergent regulation by retinoid receptors and TFs they induce. This highlights a striking potential point of convergence between genetic risk loci and an environmental risk factor for MDD.

## METHODS

### Identifying candidate psychiatric GWAS regulatory variants

To prioritize putatively regulatory variants from neuropsychiatric disease GWAS regions (predominantly MDD; **Figure 1A**), SNPs in linkage disequilibrium (LD) with GWAS tag variants at R^2^ > 0.1 were collected and intersected with histone modification, eQTL, Hi-C, and enhancer segmentation datasets from human postmortem tissue and neural lineage cell lines (see *Supplemental Methods*, **Figure 1B).** SNPs were manually selected based on diversity and density of annotation overlap within each locus (detailed in *Supplemental Methods*.) As a negative control, we identified candidates from one additional locus associated with several anthropomorphic traits^60^, in a blinded manner. Altogether, 1453 SNPs were selected. Final LD of selected SNPs was distributed similarly to starting SNPs (**Fig 1D).**^59^ Further included was a positive control TR SNP functionally demonstrated in mouse retina and brain^55^.

**Figure 1:**
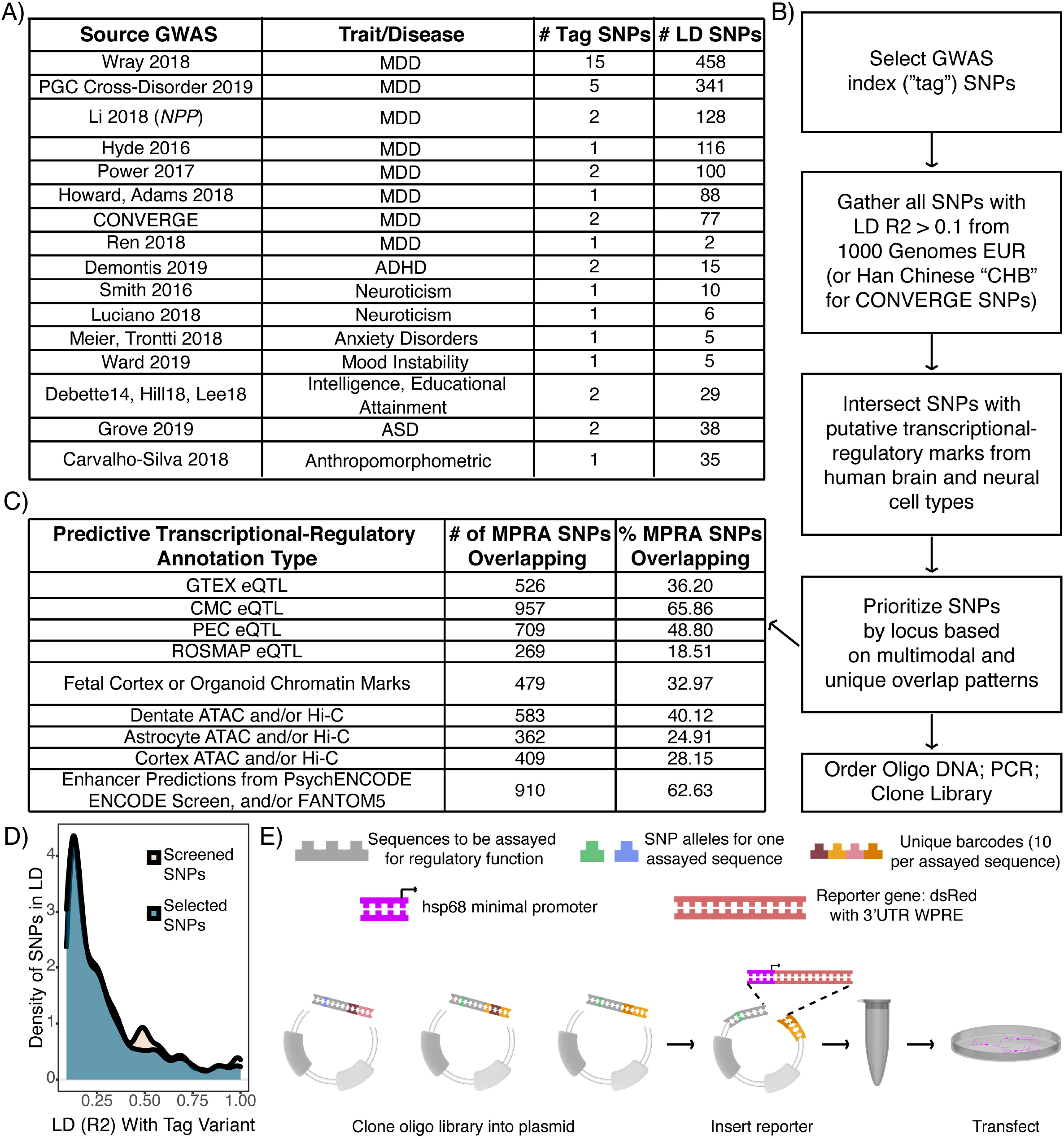
Design of an MPRA library to identify candidate functional SNPs in MDD loci. A) Table of GWAS studies and number of loci covered in the MPRA library. B) Flow chart of design and prioritization process. C) Brain and neural transcriptional-regulatory predictive annotation overlap with SNPs included in MPRA library. Fraction and number of SNPs in designed MPRA library intersecting each transcriptional-regulatory predictive annotation type. D) The manual prioritization process was not LD biased. The subset of prioritized SNPs are spread over LD space similarly to the full set of screened SNPs. E) Schematic of library construction and delivery. Panel adapted from^47^.

Human genomic sequence (hg19) tiles up to 126bp were taken centered on the 1454 candidate enhancer SNPs, each paired to 10 unique 10 base pair barcode sequences per allele and ordered as an oligonucleotide pool. Further included in the pool were 110 “basal” barcodes, where no human genomic sequence was placed upstream of the minimal promoter (i.e., such that the only variable sequence was the barcode itself). PCR-amplified oligos were then cloned into plasmid (**Fig 1E**), with subsequent insertion of a reporter cassette containing a constant minimal promoter (*hsp68*) driving the dsRed reporter gene^61^, and the untranslated “woodchuck” element (for RNA stabilization, to improve signal)^62^.

### Massively Parallel Reporter Assays

N2A cells were grown in uncoated 6-well plates in medium consisting of 0.1μM vacuum-filtered DMEM with 10% Fetal Bovine Serum (2% fetal bovine serum for the ATRA assay, based on media conditions from the literature^63,64^). For transfection, cells were reverse transfected by plating in antibiotic-free medium onto pre-plated 400μL mixtures of 2.5μg plasmid with Lipofectamine 2000. After 7 hours of incubation at 37°C and 5% CO_2_, medium was replaced with the respective medium containing antibiotics, and in the second assay, a final concentration of 20 μM ATRA dissolved in DMSO, or equivalent volume of vehicle (DMSO). Medium was not replaced before RNA collection in the first assay; in the second assay, it was refreshed every 24 hours. 72 hours after transfection, cells were collected and RNA extracted using the Zymo Clean-and-Concentrator 5 kit per manufacturer instructions. Eluted RNA was treated with Turbo DNA-free kit to remove any residual plasmid to prevent contaminating DNA reads during sequencing, and cleaned a second time using the Zymo kit as above.

### Targeted sequencing of RNA and input plasmid

Briefly, equal amounts of RNA (1μg) from each sample were prepared for sequencing by targeted cDNA synthesis using a primer against the distal 3’UTR of the reporter. These, along with input plasmid, were subjected to PCR, enzymatic digestion, ligation of Illumina sequencing adapters, and a final PCR to add sample indexes for sequencing. Enzymes, and size-selection cleanup steps used in this process are fully detailed in *Supplemental Methods.* No-reverse-transcriptase controls utilizing sample RNA were co-prepared for both experiments and did not generate detectable product, indicating sequencing amplicons generated from RNA samples were exclusively representative of RNA content. Samples were sequenced to an average depth of ~8 million reads (first assay) or ~20 million reads (second assay).

### Analysis

Allelic SNP effects on expression in the first assay and in single-condition analyses of the second assay were assessed by *t*-testing the element’s expression of each allele across replicates. dbSNP-assigned reference (“ref”) and alternative (“alt”) alleles for each SNP were used to define comparison direction (the difference of activity under the alt allele vs. the ref allele). For the first MPRA and single-condition analysis of vehicle samples from the second assay, P values were adjusted using empirical *q*-value correction via simulated allelic comparisons between random subsets of “basal” barcodes (see *Supplemental Methods*) following an analogous procedure from a multiplex CRISPR study^65^, with significance defined as q<0.05 unless specified otherwise. This ensures that a representative cross-section of expression variability driven by barcode sequences is accounted for when assessing TR differences. Single-condition analysis of ATRA samples utilized standard Benjamini-Hochberg FDR correction, as primary effects of interest in these samples were ascertained by linear modeling. For analysis of ATRA effects, barcode level expression values for both alleles of a SNP from all samples were used in a linear mixed model (LMM) with a random variable for replicate (to account for well-specific effects) as: barcode expression ~ allele + drug + allele:drug + (1|replicate). Empirical q-value correction for LMM *F* statistics was performed in an analogous manner to the prior experiment, generating a vector of F statistics for each coefficient from 20 000 randomized basal-only comparisons. For SNPs with significant allele and interaction coefficients, a meaningful allele main effect was considered present if the single-condition vehicle and ATRA analyses showed the same allelic direction of effect, with a vehicle *q* < 0.1 and ATRA FDR<0.1 (*i.e.,* near-significant within each condition of n=6, thus reasonably capable of achieving significance in the LMM analysis of the two conditions combined).

### Motifbreakr analysis and functional SNP enrichment for perturbed motifs

The motifbreakR^66^ package was used to identify TF binding motifs significantly different between alleles of each SNP. Briefly, the number of MPRA-identified functional SNPs matching a given motif for at least one allele was compared to the number of non-functional SNPs matching that motif across 10,000 random draws of *n* (number of significant) SNPs. A second version of this analysis focused on the concordance rate – that is, whether the frequency of functional variants experiencing concurrent strengthening of motif and expression or vice versa—was significant compared to 10,000 draws of *n* random SNPs from the analyzed set. In the first assay, SNPs were defined as functional based on a *q* value threshold of 0.05. In the second assay, we performed two comparisons. One analysis compared allele main effect SNPs at q<0.1 vs. those subject to neither main effects of allele or drug, nor their interaction (all q>0.1), representing the breadth of functional variant-susceptible *cis*-regulators. The second analysis compared interaction SNPs at q<0.05 vs. SNPs with an allele main effect (q_allele_<0.1) but no interaction (q_interaction_>0.1).

## RESULTS

### Many MDD loci contain more than one functional SNP

We identified >1 000 SNPs from MDD-associated GWAS loci, prioritizing SNPs overlapping with epigenetic data from neural samples, and cloned them into an MPRA library **(Figure 1)**. We included one positive control SNP, shown to alter neural tissue gene expression, and one control locus near *CDKAL1* not *a priori* associated with psychiatric disease. To identify functional variants from these SNPs, the library was transfected into N2a (also known as Neuro2a) cells (n=6 replicates, **Fig 1E**). Variant activity was assessed by RNA sequencing and barcode counts compared to input plasmid barcode counts. After filtering for read depth and barcode representation, 1013 SNPs spanning all 40 LD regions remained for analysis. Results were highly replicable across samples (Pearson *r* 0.63-0.85 for barcode expression; 0.90-0.96 for elements, **Supplemental Figure 1**). We use “element” to signify the set of barcodes corresponding to one unique sequence of interest (1 SNP = 2 elements).

Of 1013 SNPs analyzed, we identified significant allelic TR (*q*<0.05) at 76 SNPs (65 from MDD loci; 1 from the control *CDKAL1* locus) across 27 of the 40 analyzed GWAS regions, with effects ranging 0.1 to 0.63 (median 0.2) log2 fold-change (**Fig 2B**). Interestingly, the functional variant from the control locus is suggestively associated (GWAS *p* < 5•10^−6^) with *“Poisoning by analgesics, antipyretics, and antirheumatics”* in UK Biobank^67^. As this likely includes attempted suicides, the SNP was retained for analyses. The positive control SNP showed the expected higher expression of the T allele at a q-value of <0.051 (**Fig 2A**)^55^.

**Figure 2:**
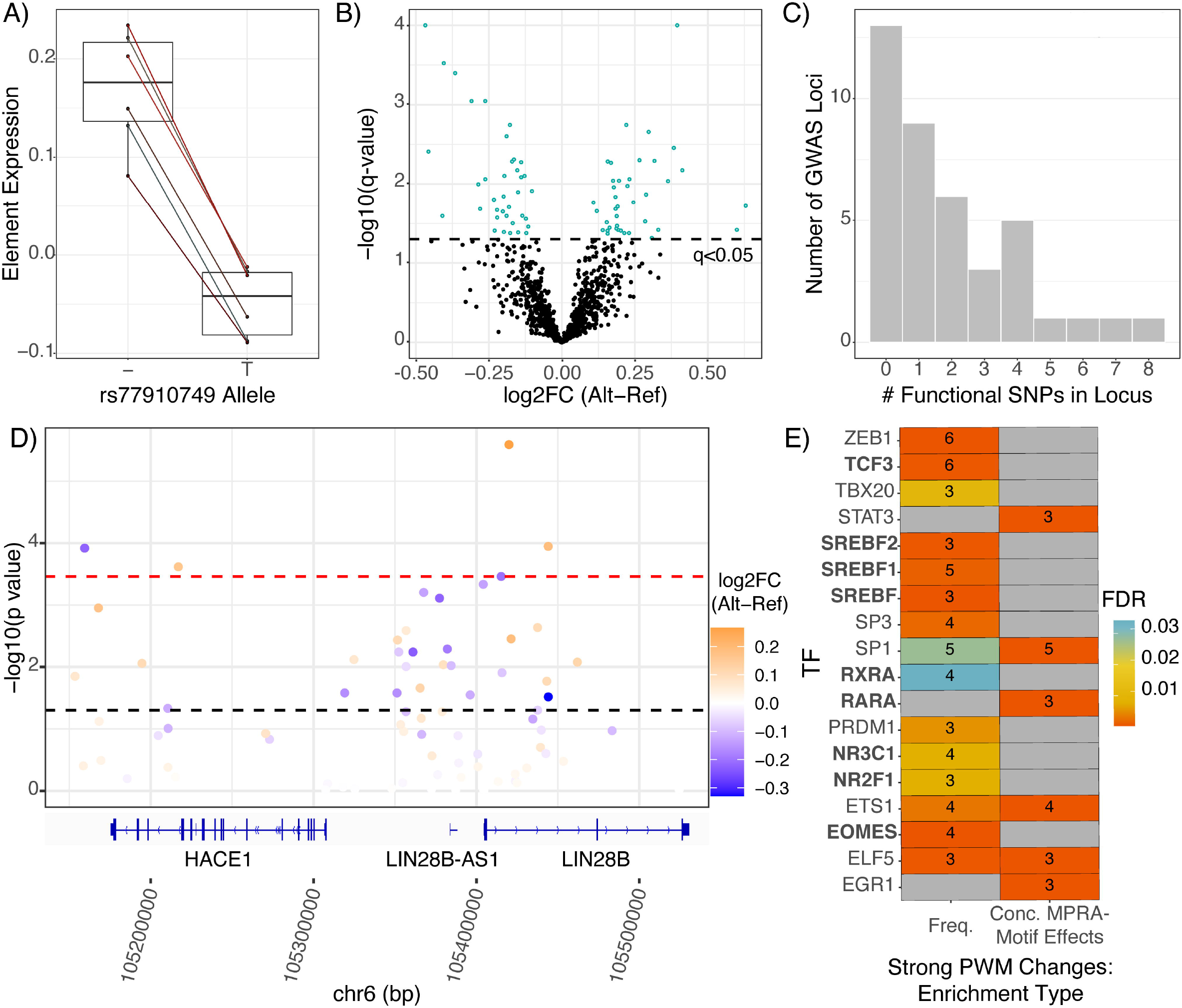
MPRA defines SNPs with a functional effect on gene expression. A) MPRA results of positive control SNP. Shen, et. al. found that the T allele drove decreased expression relative to the deletion (“-”) allele, which was robustly reproduced in the present assay^55^.) B) Volcano plot of allelic differences in reporter expression. Points represent one SNP’s composite log2 allelic fold change (alt vs. ref), determined as the mean of samplewise alternative allele barcode expression minus the matched mean of reference allele barcodes. The dotted line indicates the statistically corrected significance threshold C) Number of functional SNPs (MPRA significant SNPs) per GWAS locus in the assay. Number of loci (y-axis) containing a given number of MPRA-significant (q<0.05) SNPs (x-axis). D) The LIN28B locus harbors several functional SNPs. SNPs are plotted according to their chromosomal position (hg19) and colored based on their composite log2 allelic fold change. Refseq genes are visualized by the Integrative Genomics Viewer^95^. E) TF binding motifs involved in retinoid signaling, steroid synthesis and response, and neural activity are enriched among functional SNPs. Boxes are colored by FDR-corrected significance of enrichment for motifbreakR-defined “strong” allelic perturbations to binding motifs among functional SNPs; the number of functional SNPs perturbing (left column) and/or with concordant motif and MPRA effects (right column) are shown. Concordant effects were defined by greater MPRA expression driven by the allele better-matched to the corresponding TF motif and vice versa—the expected behavior of strictly activating TFs.

Consistent with prior studies of “conditional” or “secondary” SNP associations—wherein additional LD SNPs have associations independent of their linked, larger-effect variant^19,68,69^ ‒we identified several loci with multiple functional SNPs (**Fig 2C**) (range 1-8, mean 2.8, median 2). Notably, we identified as functional rs1806153, which was recently defined as a “conditional SNP” for MDD^18^. Our findings support models predicting multiple functional SNPs in GWAS loci, and directly validate one such finding from association analysis.

One notable TR SNP we identified, rs314267, comes from a “*LIN28B*” (nearest gene) GWAS locus repeatedly linked to MDD^9,12^ as well as cross-psychiatric disorder risk^40^. MPRA significance and effect size are illustrated for the region, showing that this locus contains several functional SNPs **(Fig 2D)**. All significant MPRA SNPs in the locus had effect directions consistent with brain eQTLs. rs314267 is the most significant *LIN28B* eQTL SNP (eSNP) in the region in PsychENCODE^70^, and is a CommonMind Consortium (CMC) eSNP for both *LIN28B* and *HACE1*^63^. *HACE1* is also downregulated in postmortem MDD hippocampal CA1^71^. Hi-C data from human neural cell cultures suggest rs314267 is within a neuron-specific *LIN28B* regulator, with promoter chromatin contacts found in dentate and cortical neurons, but not astrocytes^72^. *LIN28B* plays broad roles in neurodevelopment^73^ and has potentially sex-differentiated functions^74–77^—which, considering sex differences in MDD prevalence and severity^78,79^—makes *LIN28B* an especially interesting gene target from this locus. Finally, we examined potential upstream TR mechanisms for SNP activity usingVARAdb^80^. Query of rs314267 revealed a two order of magnitude allelic difference in the motif match p-value for *TCF4*—a gene itself implicated in cross-psychiatric-disorder risk^40,81^. Overall, the identification of functional SNPs implicated in regulation of *HACE1* and *LIN28B* exemplifies the ability of MPRAs to identify functional variants involving sensible TR mechanisms target genes.

### Shared regulatory architecture across distinct loci

We next sought to test our hypothesis that functional MDD risk variants shared retinoic acid-related TR architecture. If so, functional SNPs should disproportionately disrupt binding sites of retinoid-binding TFs compared to SNPs without an allelic effect on TR. Such data would indicate that MDD risk is mediated in part through perturbations of specific upstream transcriptional circuits, and may highlight how risk conferred through retinoids converges with risk conferred through genetics to perturb downstream gene expression.

To take an unbiased approach to our retinoid hypothesis, we first broadly analyzed all motifs showing enrichment at TR SNPs. Motifs for several dozen TFs were perturbed by the functional SNPs more frequently than expected, often with ‘strong’ perturbations to motifs and/or overrepresentation of concordant expression effects **(Fig 2E, Supplemental Figure 2)**. This included several TFs aligned with biological processes relevant to psychiatric disease. For example, several TFs are involved in steroid pathways, from regulating biogenesis (*SREBF* family, 6 SNPs) to conveying downstream TR effects——most notably, via glucocorticoid receptor (*NR3C1,* 5 SNPs; **Supplemental Figure 3**), a central component of the stress response. Functional SNP overrepresentation of SREBF motifs is consistent with high expression of these TFs in N2as and related neuroblastomas^82,83^. A second group of transcription factors included three TFs involved in neural lineage commitment/development: TCF3^84,85^, EOMES, and NR2F1^86^ (6,4, and 3 SNPs, respectively). Altogether, functional SNP enrichment for these TFs’ motifs bolster our confidence in this approach, as a) detected variation involves TFs known to be expressed in N2As (*SREBF*); b) functional variation involves TFs with roles in developing CNS, where disease variants likely act; and c) that the single-best characterized trigger of MDD (stress) is reflected in enrichment of alterations to NR3C1 motifs.

Finally, consistent with our hypothesis of convergence on retinoid-mediated TR, functional variants were enriched for “strong” perturbations of retinoid receptor TF motifs (**Fig 3**), including RARA, RARB, and RXRA (5 SNPs from 4 MDD loci, **Fig 3**). Especially notable is the motif configuration at SNP rs34416841, which falls within three partially overlapping motifs for retinoid TFs. In addition, the elements overlapping rs489591 and rs13330178 appear to be functional human retinoid TF binding sites *in vivo* based on DNAse hypersensitivity footprinting^87^.

**Figure 3:**
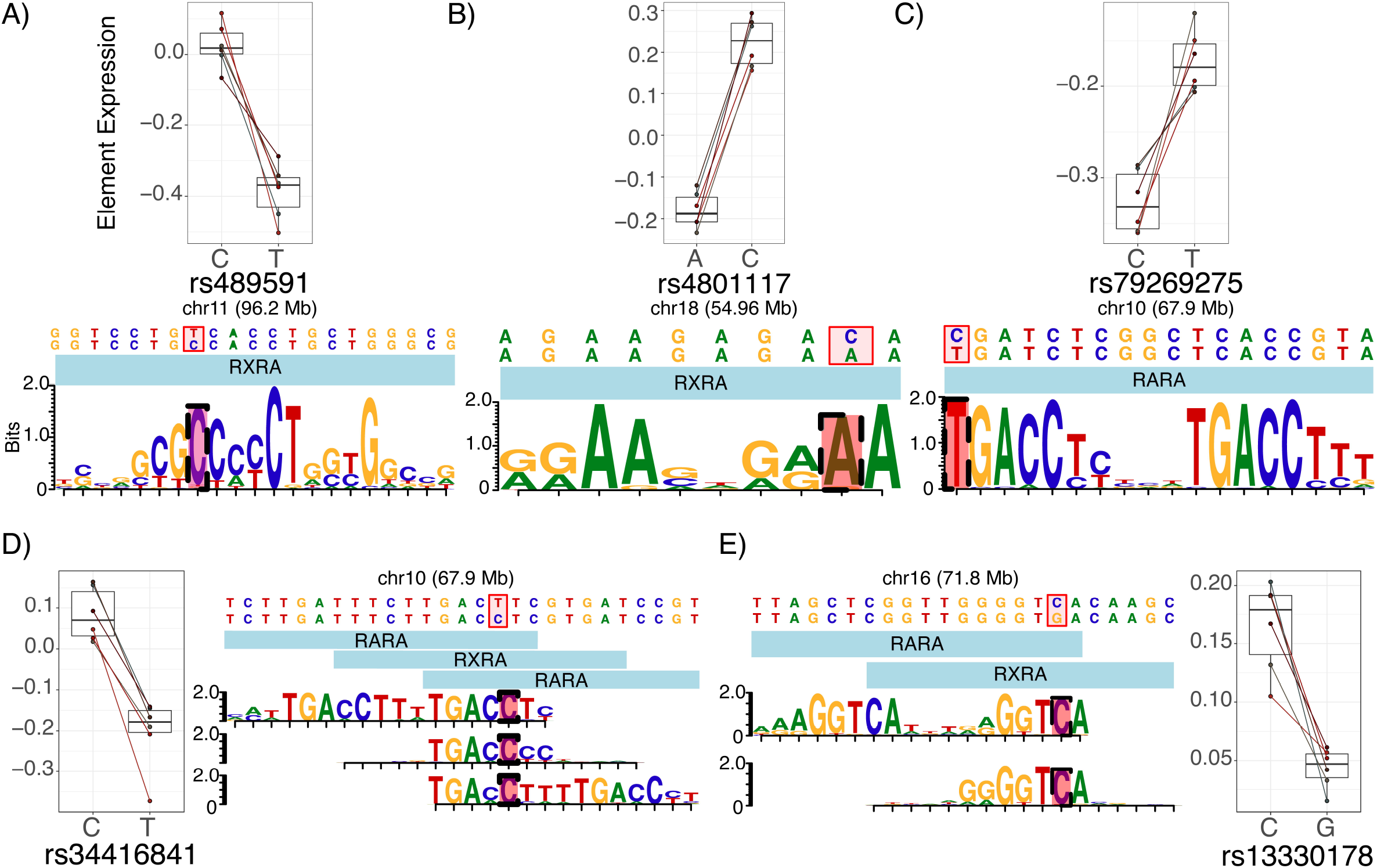
MPRA Signal at SNPs Disrupting Putative Retinoid TF Motifs. The five functional SNPs driving enrichment signals for retinoid TF motif perturbations are shown, along with the MPRA results for each variant. Each motif diagram shows only distinct position-weight matrices (PWMs). (motifbreakR uses a large meta-collection of motifs, which were often identical or nearly identical across retinoid TFs; such redundant motifs are not shown). Among functional-SNP enriched retinoid TFs, A) rs489591 and B) rs4801117 exclusively perturb RXRA motifs. C) rs79269275 perturbs an RARA (or RARB, identical but not shown) motif. D) rs34416841 alters several similar retinoid motifs across multiple positions and TFs. Not shown: near-identical motifs for RARB along the same sequence as the 5’ RARA motif; near-identical RXRB and RARB motifs along the same sequence as the center RXRA motif; and a near-identical RARB motif along the same sequence as the 3’ RARA motif. E) rs13330178 disrupts an RARA or RXRA binding site. Given that RXRA and RARs are known to heterodimerize, it is possible that this SNP disrupts the RXRA component of such a heteromer’s binding sequence.

### Retinoids unmask additional functional SNPs in MDD loci

Our findings supported the hypothesis that MDD-associated variants across multiple loci converge on TR that may be modulated by retinoids. We thus designed a pharmacological follow-up with two goals in mind. The first goal was to functionally verify that retinoids were involved in TR at SNPs where their motifs were found (in *cis*), and potentially unmask additional retinoid-targeted alleles. Our second goal was to further assess retinoid signaling *trans* (i.e., indirect) effects on variants from these same GWAS regions, *e.g.,* via non-retinoid TF induction, co-regulation, or repression^32^. Therefore, we performed a second MPRA adding an all-*trans* retinoic acid (ATRA) condition.

After 48 hours, cultures were imaged to verify drug delivery and activity (as ATRA is light-sensitive) based on known morphologic responses of N2as to ATRA, which include neurite outgrowth and mitotic arrest^58,88,89^. Indeed, drug-treated cells had a qualitatively lower cell density and produced neurite-like processes (**Fig 4A**) in comparison to vehicle-treated cells **(Fig 4B**). After RNA sequencing, we first analyzed vehicle-treated replicates alone to ensure replicability of the assay. Element expression levels in the vehicle condition strongly correlated to the first experiment (Pearson *r* = 0.91; **Fig 4C),** and replicated the functional variants (**Fig 4D**); all 31 shared significant SNPs showed consistent directions of effect.

**Figure 4:**
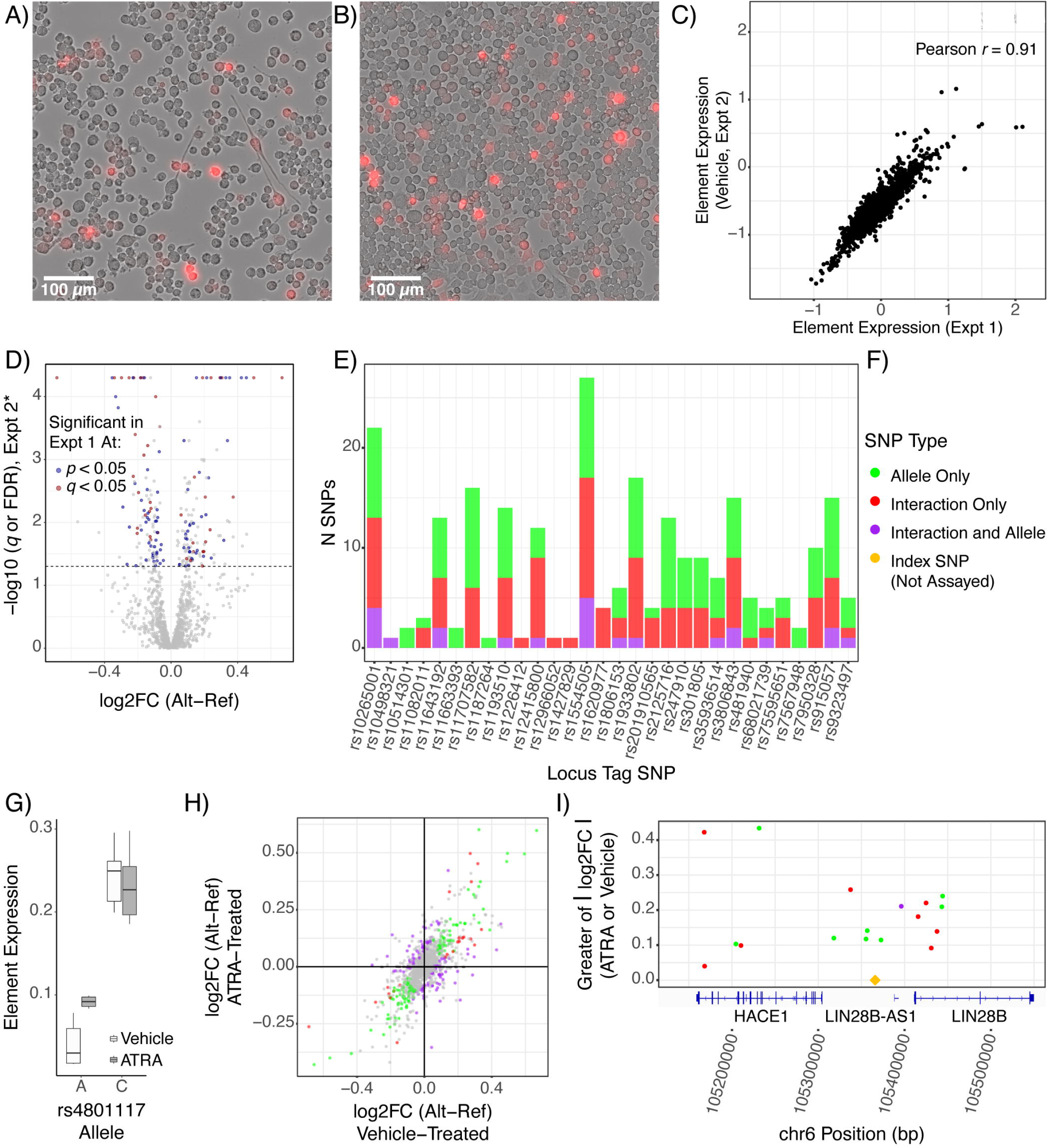
Retinoid treatment alters transcriptional regulation and unmasks additional functional variants. A) ATRA-treated cells show growth arrest and neurite growth, demonstrating effective ATRA treatment, while B) Vehicle-treated cells continued to proliferate in a de-differentiated state. C) Results of the vehicle treatment replicate the initial MPRA findings. Element (single-allele) expression values for each sequence assessed in both assays is plotted. D) Significant and marginally significant functional SNPs from the first assay showed effects in the second assay. The larger allelic difference value from the ATRA and vehicle single-condition analyses is plotted for each SNP on the x-axis; the y-axis value is the corresponding corrected p-value (FDR correction for the ATRA-only analysis or empirical q-value correction for vehicle-only analysis). E) Retinoids unmask functional SNPs with additional or exclusive retinoid-mediated effects. F) SNP effect(s) color key for panels E, H, and I. SNPs with both effects were those with LMM interaction q<0.05, LMM allele q<0.05, and both single-condition analyses showing the same allelic effect directionality at ATRA FDR<0.1 *AND* vehicle q<0.1. F) rs4801117-A shows greater activity with ATRA treatment while the G allele is unaffected. The ATRA having an expression effect only on the A allele is consistent with the A allele matching the RXRA motif as shown in Fig 3B. H) Transcriptional-regulatory SNPs show a wide range of altered and unaltered effects with ATRA treatment. Single-condition log2FC values are shown. I) Several additional SNPs with retinoid-dependent function (i.e., allele-ATRA interaction) in the *LIN28B* locus. Only significant SNPs are illustrated. Notably, there are several functional SNPs clustered around the GWAS index SNP, suggesting association signal at this locus may be driven by multiple functional/conditionally functional variants.

We next applied a linear mixed model (LMM) to identify SNPs responding to ATRA (that is, allele-drug interactions). A total of 1079 SNPs were analyzed after filtering for read and barcode depth. In part due to the effective doubling in power to detect allelic effects with 12 replicates and the LMM approach, we now identified 137 variants with a main effect of allele (129 from MDD loci). Four of the five retinoid receptor motif-perturbing variants from the first assay passed filtering; all four of these variants again showed allelic main effects (all *q* < 0.01), as did many other functional variants identified in the previous experiment **(Fig 4D**). To our surprise, more variants showed a significant drug-allele interaction effect: a total of 128 SNPs (122 from MDD loci) (**Fig 4E-F**). Among the drug-allele interaction SNPs were one of the four retinoid-related SNPs identified from the first assay (rs4801117; q_interaction_<0.025, **Fig 4G**), while another trended towards interaction (rs489591; q_interaction_=0.117). This strongly supports a role of retinoid TF activity at rs4801117 as predicted by the motif analysis. More broadly, comparison of changes between the two conditions reveals the striking extent to which the regulatory landscape of the N2As was altered by ATRA (**Fig 4F and 4H**). Notably, several additional functional variants were identified in the previously highlighted *LIN28B* locus, further illustrating multi-variant and context-dependent aspects of GWAS loci (**Fig 4E and 4I).** In all, this experiment highlights the ability of MPRAs to detect contextual influences such as cell states and signaling on functional noncoding variation, and to unmask distinct, context-dependent functional SNPs.

### Retinoids reveal additional axes of convergent regulation at functional MDD-associated SNPs at levels of TF and cell type

As the ATRA-based assay provided improved power to identify allelic variant effects on expression, we sought to again assess convergent transcriptional effects of these regulatory variants by employing the same motifbreakR-based analyses. When examining SNPs with allelic effects in comparison to SNPs with no allelic, drug, or interaction effects, several retinoid receptor motifs were again overrepresented, including those of RXRA, RXRB, RARA, and RARG **(Fig 5A, Supplemental Figure S3)**, totaling 11 of the 92 allele-main effect SNPs analyzed and spanning 10 MDD GWAS loci. These findings further support retinoid receptor binding sites as an upstream regulatory system recurrently involved in MDD risk genetics.

**Figure 5:**
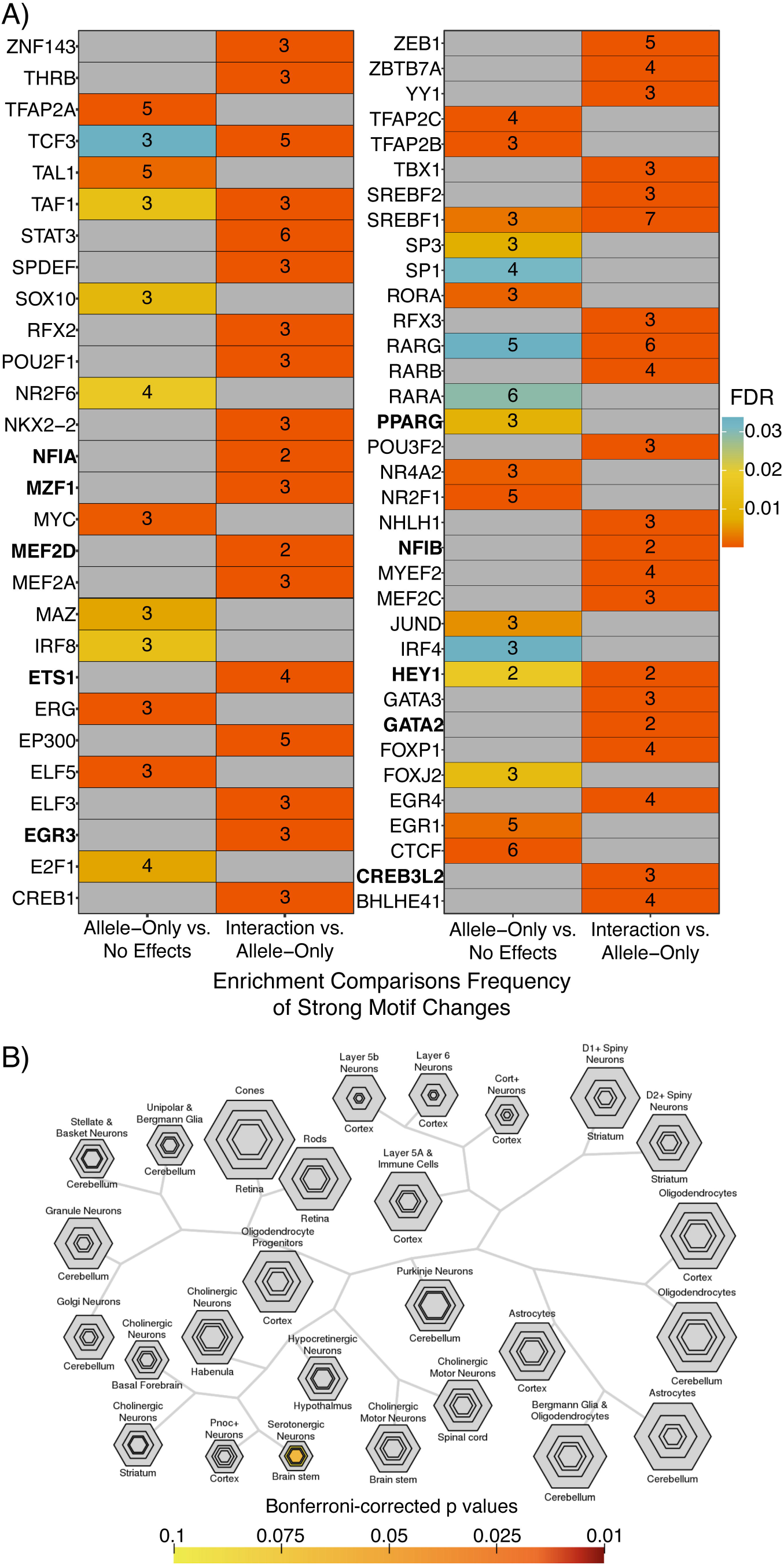
Distinct TFs underlying retinoid-dependent functional SNPs and implication of serotonergic neurons. A) Motifs overrepresented among ATRA-independent (allele effect without interaction) SNPs (left columns) or among ATRA-dependent (interaction) SNPs (right column). The heatmap is shown in halves for visibility. TFs identified as ATRA-upregulated in human neuroblastoma lines^88^ are in bold font. B) TFs implicated by ATRA-interacting SNPs significantly overlap TFs enriched in serotonergic neurons. Plot generated using the cell-specific expression analysis (CSEA) tool (http://genetics.wustl.edu/jdlab/csea-tool-2/)^31^.

As retinoids resulted in stark changes across the transcriptional-regulatory landscape, we further sought to understand the TFs potentially underlying allelic effects with retinoid exposure. Therefore, we also analyzed the interaction SNPs in comparison to allelic SNPs that were *not* subject to interactions. This revealed a novel set of TFs not observed in the preceding analyses, including TFs with roles in neural differentiation and maturation (**Fig 5A, Supplemental Figure S3)**. We compared the overrepresented motifs to TFs recently demonstrated to be upregulated in neuroblastoma lines (KCNR, LAN5) by ATRA. Of the 26 TFs identified as ATRA-induced in these two lines, motifs were available for 18 in our analysis. Of these, motifs for 6 of the TFs were enriched among allele main-effect-only SNPs, while motifs for 12 of these TFs were enriched among the retinoid-allele interaction variants^88^ (**Fig. 5A, Supplemental Figure S3**), supporting our predictions of TFs playing ATRA-dependent roles at functional SNPs.

Comparison between ATRA-interacting and -independent functional SNPs results should properly control for motif matches that do not correspond to retinoid-bound TFs (as ATRA agonism of retinoid receptors should modify their activity). Indeed, in contrast to our preceding motif analyses of allelic effect variants, only two retinoid receptors were enriched among the retinoid-interacting functional variants: RARG (eight SNPs, five of which also matched RARB motifs) (**Fig 5A, Supplemental Figure S3)**. This suggests that RARG and/or RARB is the particular retinoid receptor at the nexus of allele-ATRA interaction effects in this model, and may likewise be TFs responsible for ATRA-dependent allelic effects at MDD risk loci overall. Altogether, these findings support cross-locus involvement of retinoid receptor binding sites—including those of RARG—and of retinoid-regulated TFs in MDD-associated regulatory variants.

Finally, we utilized these sets of TFs as gene sets to investigate whether retinoid-dependent or -independent regulatory variants might be particularly active in certain cell types of the brain. Using a collection of previously published translatome-wide brain cell type-specificity metrics^31^, we screened these TFs for enriched expression in specific cell types. Three TFs (spanning 8 ATRA-interacting SNPs) were discovered to be highly specific to serotonin neurons (**Fig 5B**): *GATA2, GATA3,* and *FEV*, while no cell type enrichments under FDR<0.1 were noted for TFs linked to ATRA-independent variants. We believe this is not an artifact of the N2a system because we could identify no evidence in the literature suggesting a serotonin-like identity of N2a cells with or without ATRA treatment. However, exogenous retinoids have been shown to lower circulating serotonin in humans^90^ and to alter morphology of rat raphe neurons in slice culture^91^, suggesting they are retinoid responsive. Single-cell RNA sequencing data from mouse brain reveals that these TFs are only expressed in serotonergic, noradrenergic, peripheral autonomic, and midbrain inhibitory neurons—with all three expressed in serotonin neurons^92^—suggesting that these cell types in particular may be a cellular point of convergence for several retinoid-mediated functional SNP effects on MDD risk.

## DISCUSSION

To date, most functional investigations of SNPs in the context of psychiatric disorders have taken place in a low-throughput manner, such as single-variant classical reporter assays^20^or using CRISPR-Cas9 technology to edit limited positions for deep phenotyping^93^. Here, we leveraged MPRA to screen over 1 000 SNPs from loci associated with MDD, related phenotypes, and broader psychiatric disease, demonstrating the utility of this technique for dissecting the functional regulatory architecture of psychiatric GWAS loci, and defining shared upstream regulatory features across loci.

In doing so, we identify over 100 SNPs with allelic effects on expression, with most coming from loci containing ≥ 2 functional SNPs. These data provide experimental support for the assumptions of polygenic/omnigenic theories, which predict multiple SNPs with allelic effects within GWAS loci. Further, we examined convergence across loci, predicted by omnigenic theory. By examining the shared regulatory features (transcription factor binding site motifs) based on enrichment at functional SNPs, we were able to predict several TFs with TR activity recurrently altered across MDD-associated SNPs, highlighting retinoid receptor TFs in particular.

Retinoids are encountered both exogenously (e.g., as ATRA in oncology, and as isotretinoin, carrying a black-box warning for suicidality) and endogenously, including during brain development. To investigate how SNP functions may be altered by retinoids we repeated the assay with an ATRA drug condition. ATRA drastically rearranged the TR landscape of N2a cells, resulting in altered and novel allelic effects at over 100 SNPs and revealing ATRA-dependent mechanisms of function across 122 SNPs from 22 of 26 MDD GWAS loci assessed. Motif-SNP perturbation enrichment identified two retinoid receptors, RARG and RARB, as the likely TFs involved in functional SNP activity at retinoid receptor binding sites in this system. This could be because either a) in N2as, RARG/RARB are the predominant retinoid receptor, or b) that these cells express a broader array of retinoid-activated TFs, but only the RARG and RARB motifs are a high-fidelity representation of “true” retinoid receptor family-wide binding sites. Nonetheless, these findings suggest that RARB, and RARG in particular, merit mechanistic follow-up regarding TR differences at MDD-associated SNPs. Future work may be able to leverage biobank-level datasets to ascertain whether retinoid-interacting SNPs are overrepresented in retinoid-treated patients experiencing adverse psychiatric side effects. While data on endogenous retinoids, *e.g.* plasma values, are not currently available in large genotype-phenotype-health record cohorts like UK Biobank, future datasets may enable investigation of circulating retinoids and their interaction with genotype in cognitive and psychiatric phenotypes.

The methodologic requirements of high-throughput assays, such as MPRAs, bring inherent limitations to their results. The primary precaution to be taken is with regard to cell type relevance. MPRAs are subject to the TR landscape of the cell type used. The cell type here was selected for its intact retinoid signaling, but is not likely central to MDD, as neuroblastomas are derived from peripheral neural crest progenitors (though they can be differentiated into dopaminergic neurons^89^, and commit to neuronal differentiation with ATRA^58,64^). On the other hand, the neural crest-derived autonomic nervous system has received little consideration (i.e., relative to brain) in psychiatric genetics of MDD despite the well-appreciated role of stress in depression. Some of the functional SNPs identified may thus play a role in the autonomic system as opposed—or in addition—to the brain, though this assay is not well designed to empirically address this, as SNPs were prioritized for assay based on brain epigenomics.

We find that principles of the omnigenic model appear to hold true for MDD risk genetics, including the presence of far more functional variants (a total of 351 SNPs with allelic and/or interaction effects of 1178 assessed across the two assays) than there are GWAS loci per se. We find, interestingly, that functional SNPs form convergent subsets of upstream (transcription-regulatory) sequences and systems, which in turn have shared retinoid dependence and are collectively enriched in serotonin neurons via eight of 113 SNPs spanning functional variants in binding motifs of TFs GATA2, GATA3, and FEV. It has previously been demonstrated that systemic administration of all-*trans* retinoic acid depletes serotonin by over 40% in the rat brain^94^, supporting the serotonin system as a convergent target of retinoid-regulated pathways. As GWAS of MDD begins to explore severe, treatment-refractory cases^7^, it will be interesting to see whether associated variation still shows such convergence, as treatment-resistant depression (generally, non-response to two or more classes of antidepressant) effectively signifies non-response to multiple serotonergic agents.

In all, we assessed the architecture of *cis-*regulatory variation in psychiatric disease risk loci, identifying at least one functional SNP in the majority of the 40 GWAS loci examined, largely corresponding to MDD-associated SNPs. Strikingly, retinoid receptor binding sites and TR systems subject to regulation by ATRA have a substantial impact on whether and how MDD-associated SNPs are functional. These findings constitute a robust experimental demonstration of the influence of physiological and environmental states on the molecular activities of disease-associated SNPs, and constitute a high-confidence set of MDD SNPs meriting deeper functional characterization of both their TR mechanisms and their environmental interactions.

## Supporting information

Table S1-All Significant Variants in The MPRAs and Each Significant Effect.

## ACKNOWLEDGEMENTS

We thank Stephen Plassmeyer and Tomás Lagunas, Jr. for technical assistance; Barak Cohen, PhD, Michael A. White, PhD, Dana King, PhD, Brett Maricque, PhD, and Ryan Friedman for library design, cloning, and analysis advice; and Genome Technology Access Center@McDonnell Genome Institute (GTAC@MGI) and Jessica Hoisington-Lopez and MariaLynn Crosby from the DNA Sequencing Innovation Lab at The Edison Family Center for Genome Sciences and Systems Biology for sequencing support. This work was supported by the NIH (1F30MH1116654 to BJM, and 1R01MH116999 to JDD) and The Simons Foundation (571009 to JDD). GTAC@MGI is supported by UL1 TR002345. SNP annotation data resources are detailed in the Supplemental Text. We would also like to thank Idoya Lahortiga, Ph.D. and Luk Cox, Ph.D. curators of Somersault1824 (https://gumroad.com/somersault1824), for their open-access, Creative Commons BY-NC-SA 4.0 licensed libraries of high-quality biomedicine graphics (especially those from Graphite Life Explorer, ePMV, and Eyewire), adapted for Figure 1E. Color palettes for plots are from the R package wesanderson (https://github.com/karthik/wesanderson).

## Data and code availability

A summary spreadsheet of all significant SNPs identified in one or both assays, along with full analysis results, including barcode-wise expression in each sample, single-condition allelic effect tests, linear modeling results, and significantly enriched TFs in each of the comparisons executed, along with the code utilized to execute these analyses, is available https://bitbucket.org/jdlabteam/n2a_atra_mdd_mpra_paper/src. Raw sequencing files are deposited in GEO under accession GSE167519.

