## Supplemental Methods and Figure Legends for "Transcriptional-Regulatory Convergence Across Functional MDD Risk Variants Identified by Massively Parallel Reporter Assays"

**Library Design and SNP selection**

The library was designed by selecting tag SNPs of interest from neuropsychiatric trait and disease GWAS studies, predominantly for MDD or multi-diagnosis groups including MDD cohorts^1–9^, and from GWAS of traits with high SNP co-heritability with MDD—namely, neuroticism^10,11^ and mood instability^12^. Additional variants discovered in other psychiatric disorder GWASes at or near MDD tag variants were included from studies of ASD^9^, anxiety disorders^13^, and attention-deficit hyperactivity disorder^14^. Two tag SNPs associated with intelligence^15^ and educational attainment^16^ near the gene *PTGER3*, which we previously illustrated to be sex-differentially expressed and functional in the mouse locus coeruleus^17^, were included. One negative control tag, rs1883640, close to the transcription start of gene *CDKAL1* and associated with several anthropomorphic traits^18^, was also included as a negative control locus. In all, 38 tag SNPs were selected for LD expansion and epigenomic overlap screening in the LD neighborhood. All LD partners at R2 > 0.65 were included without consideration of epigenomic annotation intersections for two additional MDD-associated tag SNPs near sex-differentially expressed genes of mouse LC—*Slc6a15* and *Lin28b*^17^; SNPs from these two loci with R2 < 0.65 were also included solely on the basis of a RegulomeDB score of ≥ 4, signifying a SNP overlapping both a TF binding site and a DNAse hypersensitive site across epigenomic data curated by the tool^19^.

LDlink (using dbsnp 151) was queried for each tag SNP to acquire all biallelic EUR (or in the case of tags from Han Chinese CONVERGE^2^, Han Chinese (“CHB”)) SNPs in LD with the tag^20^. These SNPs were subsetted to those with a minor allele frequency ≥ 1% and with an LD R2 > 0.1 with the tag, as GWAS studies generally define ‘independent loci’ as SNPs in with LD R2 < 0.1. The hg19 coordinates of the retrieved, and subsetted LD SNPs were then intersected to those of myriad CNS epigenomic datasets (also in hg19 coordinates), including postmortem brain tissue eQTLs from GTEX v7^21^, PsychENCODE^22^, the CommonMind Consortium^23^, the Lieber Institute/Brainseq Consortium^24,25^, and ROSMap^26^; enhancers predicted based on human postmortem adult and fetal brain tissue histone marks or enhancer RNAs^22,27–31^; and chromatin contacts for human neural cell types identified *in vitro*^32^.

LD SNPs were selected by manual inspection of intersecting epigenomic annotations within an LD block, including the negative control *CDKAL1* block. Negative control locus SNPs were selected while blinded to control status of the locus (by replacing the parent locus’ name as one coming from a sub-region of interest during the selection process). 11 SNPs were removed from consideration due to overlap with a nonsynonymous coding SNP. For inclusion, a SNP was required to be an eQTL in at least one of the brain datasets mentioned and intersect with ≥1 additional annotation track; two exceptions to this were that no overlaps were required for the SNPs included by RegulomeDB score (see above), and a single overlap (eQTL or otherwise) was considered adequate for inclusion in a small minority of loci where most SNPs did not intersect any annotation. In annotation-rich regions, SNPs with the greatest diversity and abundance of intersecting annotations were selected. While computationally selecting SNPs based on the greatest number of intersections would be a more time-efficient approach, recent work suggests that regulatory variants may be better predicted by training deep learning models on local genomic sequences rather than sequences from across the full genome^33^.

To design the MPRA oligonucleotide sequences ordered, genomic tiles of 126bp, centered on the variant of interest were extracted from human reference genome hg19. For variants where the alleles were not of the same length (e.g., single-base deletions or multi-base alleles), the longer allele’s sequence spanned 126bp, with the shorter allele spanning 126-(difference in length) bp. As oligonucleotide synthesis requires uniform sizing, all oligonucleotides were brought to a final length of 200bp by adding bases between cut sites used for directional insertion of the reporter gene, such that the bases are absent from the final plasmid library.

Sequences containing cut sites that would interfere with cloning were taken from the smallest up- or down-stream window to keep the SNP as near the center of the sequence as possible, by shifting enough to trim away 1-2 bases of the interfering cut site. When this required shifting sequences more than ±30 bp, the selected SNP was removed from the design process (resulting in the removal of 8 SNPs from the selected set). The final, cut-site-free genomic tiles representing each allele of the 1453 selected SNPs were programmatically added to random ten, 10bp barcodes, which were pre-subsetted to be: ≥ 2 Hamming distances from one another, 25-75% GC, without >4 of any one nucleotide in a row, and without restriction sites or partial restriction sites that would recreate a full site during the cloning process. Control sequences were likewise paired to 10 barcodes each. Finally, ‘basal’ oligonucleotides with 126 bases filling the space between gene-insertion cut sites (to ultimately place the minimal promoter directly adjacent to the reporter gene) were generated and paired to 110 barcodes to get an accurate measure of transcriptional output driven by the hsp68 minimal promoter in the absence of upstream mammalian genomic sequence.

**Cell culture, transfection, and RNA collection.**

Mouse neuroblastoma N2A cells were grown in uncoated 6-well plates in medium consisting of DMEM and 10% Fetal Bovine Serum (2% fetal bovine serum for the ATRA assay, based on media conditions from the literature)^34,35^. Cells were fed 2mL per well at each passage, and medium was refreshed as needed indicated by yellowing of the phenol red indicator contained in the DMEM solution. Cells were incubated at 37°C, 5% CO2.

For transfection, a tube containing 10µL per replicate of Lipofectamine 2000 in 200µL per replicate OptiMEM was prepared and incubated at room temperature for 5 minutes. Meanwhile, a second tube containing 2.5µg of MPRA plasmid library per replicate and 200µL of OptiMEM was prepared. The contents of the latter were added to the tube containing Lipofectamine 2000 and incubated at room temperature for 30 mintues. The mixture was then aliquoted out in volumes of 400µL to each well (replicate) and the plate re-lidded. The plates stayed 45-90 minutes in the biosafety cabinet until cells, grown and split from the same parent culture in both assays and collected from all wells of a well plate (first assay) or T150 culture flask (second assay), were aliquoted on top of the 400µL of plasmid•lipofectamine•optiMEM mixture.

Cells to be transfected were lifted from their existing wells by adding 1mL of 0.25% Trypsin•EDTA and incubating at 37°C for 5 minutes. 2mL of DMEM were added to each well, and the full 3mL of DMEM, trypsin, and cells were collected from each well into a tube containing *n* mL of 10% FBS in DMEM, where *n* was the number of wells being passaged. Cells were spun down at 700 rcf for 5 minutes at room temperature, then resuspended 10mL in antibiotic-free N2A medium (for initial MPRA, 0.1µM vacuum filter sterilized 10% FBS in DMEM; for retinoic acid MPRA, 0.1µM vacuum filter sterilized 2% FBS in DMEM). Duplicate counts of cells were taken using the Countess automated cell counter. The average number of reported viable cells per mL was used to determine plating volumes to achieve target cell densities in 2mL volumes, and the required volume of the same medium used for resuspension (less 10mL) was added to the resuspended cells before aliquoting. For the initial MPRA, these target densities were three 9.6 cm^2 wells with 5.7•10^5 cells/well and three 9.6 cm^2 wells with 8.3*10^5 cells/well. Linear modeling of results from the first assay revealed singular fits when attempting to fit a variable for initial plating density, indicating this played no role in the detected expression. In the retinoic acid MPRA, twelve 9.6 cm^2 wells at 7.8*10^5 cells/well. 2mL of resuspended cells were aliquoted to each well, then incubated at 37°C for (6.5 hours in initial MPRA, 7.5 hours in retinoid MPRA). At that time, medium was removed and replaced with the respective experiment’s N2A medium, now containing antibiotics (and in the case of the retinoid experiment, either 20µM all-trans retinoic acid in DMSO, or an equivalent amount of DMSO alone for vehicle). For the initial MPRA, medium was not changed before cells were collected; for the retinoid MPRA, medium was changed every 24 hours with a freshly prepared aliquot of N2A medium supplemented with respective drug or vehicle. Retinoic acid and vehicle were prepared in a dark room in 500 and 200 µL aliquots, respectively, stored in black microfuge tubes at -20°C and handled opened only in biosafety cabinets without room or hood lights on.

In the initial MPRA, cells were collected 72 hours after transfection by rinsing wells twice with 1mL of DPBS, then adding 1mL of DPBS, thoroughly lifting cells with a scraper, and collecting the full 1mL of cells from the well into its own microfuge tube. Cells were spun down at 3000 rcf at 4°C for 10 minutes. 750µL of supernatant was removed, and replaced with 750µL Trizol. Samples were shaken vigorously by hand for 10 seconds to lyse cells and incubated on the benchtop at room temperature for 10 minutes. In the retinoic acid MPRA, the same rinses were performed; followed by adding 250µL dPBS, then 750µL of Trizol LS to each well. The plate was swirled until cells were clearly lysed, at which point the full 1mL volume of each well was collected into a centrifuge tube, allowed to stand at room temperature for 10 minutes.

The handling of samples from both experiments was the same from that point forward: 200µL chloroform was added to each tube, which was then shaken vigorously by hand for 30 seconds. Tubes sat at room temperature for 7 minutes, and then were spun down at 12,000 rcf at 4°C for 30-40 minutes. 325µL of the aqueous phase was collected, and added to tubes pre-filled with two volumes (650µL) of Zymo RNA Binding Buffer and their combined volume (975µL) of 100% ethanol. The RNA solution was then purified using the Zymo Clean and Concentrator-5 kit per the manufacturer directions. RNA was eluted into 40µL of nuclease-free water.

To remove any residual plasmid from the samples, the Turbo DNA-Free kit was used to perform vigorous DNAse treatment (2µL of DNAse per RNA sample) according to the manufacturer instructions. RNA with DNAse in solution was incubated at 37°C for 1 hour, followed by addition of the kit’s DNAse inactivating matrix and a 10 minute spin at 8000 rcf at room temperature. 44µL of supernatant was collected, and subjected to a second round of purification using the Zymo Clean and Concentrator-5 kit as described above. (A second purification was required in order to remove an unknown component of the Turbo kit which otherwise interfered with RNA concentration/integrity measurements).

**Sequencing library preparation.**

1µg of RNA was subjected to target-specific cDNA amplification using a primer reverse complementary to the proximal polyA signal sequence in the reporter transcript. 20µL reactions containing 1µg RNA, 4µL 5x SSRT first strand buffer, 0.6µL rRNAsin (Promega), 0.9µL SSRT III enzyme, 1µL 12µM antisense primer, 1µL 100 mM DTT, and 1µL 10mM dNTPs, brought to volume with water. Reactions were incubated at 50°C for 1 hour. Remaining enzyme, primers and RNA were then removed by adding 3.5µL Exosap-IT and incubating for 15 minutes at 37°C, followed by addition of 1µL 0.5M EDTA, addition of 3µL 1M NaOH with heating at 70°C for 12 minutes, and neutralization with 3µL 1M HCl.

The single stranded cDNA was then purified by washing 10µL MyOne silane beads per sample with Buffer RLT (Qiagen), resuspending beads in 3•sample volume of RLT, and adding to each sample, mixing end-over-end for 30 minutes to bind cDNA. Beads were magnetized and washed twice with 80% ethanol, air dried for 5 minutes, and bound cDNA eluted into 12µL 5mM Tris HCl pH 8. 10µL of this product (cDNA) or 80ng transfected plasmid DNA were then used to generate double-stranded, target-specific product from the cDNA using the same antisense primer as reverse primer and a primer in the 3’UTR WPRE element of the reporter as the forward primer for 15 cycles with Phusion HF 2x Master Mix. Product was size-selected using Ampure XP beads by bringing the 20µL PCR reaction to 100µL with water, adding 80µL beads, magnetization, capture of 160µL supernatant and adding 40µL fresh beads to it, magnetization, 3x washes with 80% ethanol, and air drying, followed by elution into 12µL 5 mM Tris HCl Ph 8. (This procedure is henceforth referenced as “80/40 Ampure XP cleanup”).

The double stranded product was then digested with HindIII-HF (NEB) and NheI-HF (NEB) in CutSmart buffer at 37°C for 1 hour, to create sticky ends flanking the barcode sequences to which sequencing adapters could be attached. The digested product was purified as above, but using 100µL beads and 50µL beads, with 180µL supernatant recovery added to the second (50µL) aliquots of beads. Product was eluted as above, using 10µL eluate for adapter ligation.

Adapters were ligated using Enzymatics T4 ligase (1µL), Enzymatics T4 10x buffer (2µL), 1µM HindIII-compatible, staggered-length (to improve read heterogeneity for sequencer compatibility) Illumina adapters for read 1 and 1µM single-length NheI-compatible adapter for read 2. Ligation ‘cycling’ was used, alternating cycles of 30s at 25°C (annealing) and 30s at 16°C (ligase activity) for 560 cycles, a procedure previously shown to improve ligation yields twofold in restriction cloning^36^. Product was purified and eluted with the 80/40 Ampure XP cleanup; 10µL eluate was used for 50µL index PCRs with sample-specific read 1 indices, user-specific read 2 indices, 2x Q5 Ultra II Mastermix (NEB) for 9 cycles. Product was purified using 80/40 Ampure XP cleanup, eluted into 19µL of 5mM Tris HCl pH 8. 17µL were recovered (2 used for verifying proper product at ~330bp was produced by using the D1000 kit with the Tapestation 4200 system). Up to 15µL of each library was mixed at an equimolar amount, with approximately 1.5 times the molar amount of the sample corresponding to input DNA for maximally accurate quantification of this single DNA sample.

**MPRA Analysis.**

Raw fastq files for each RNA sample are initially processed by using the bash terminal and the string-matching command, sed. The string matching requires an exact match of 6bp upstream sequence and 8bp downstream sequence corresponding to the restriction-ligation sites wherein the reporter was ligated to the barcode, and the barcode to the plasmid, respectively. For each string matching this pattern exactly, the 10 bases of barcode sequence between are extracted and put in a text file for the corresponding fastq. To then get barcode counts, sequences in the text file of extracted barcode-positioned 10mers are subsetted to those containing a perfect 10 base match to a barcode sequence in the library and are counted using the R function *table* , which tabulates the occurrences of each barcode sequence specified in a reference metadata file (containing the designed barcodes only and related information) for each sample’s string-extraction output /text file. This process is extraordinarily efficient, requiring only 1-2 minutes per 20 million reads on a personal computer.

Following the tabulation of barcode counts, barcode counts are converted to counts per million (mapped) before filtering. Several filtering steps are then used to ensure robust representation of barcodes, their paired sequences, and samples before testing expression effects. These steps proceed in the following order. 1) Barcodes with a DNA count under a specified threshold (75) are struck from the counts table from the DNA *and* RNA samples. 2) For sequences with 4 or fewer remaining DNA barcodes represented across all samples after this step, all other barcodes for the sequence are struck from the all samples in the table to avoid analysis of sequences with inadequate barcoding depth. 3) Counts of RNA barcodes are then removed on a per-barcode-per-sample basis if they fall below a separate minimum read threshold (30 counts), set below the DNA threshold to allow for detection of repressive effects). At this point, preliminary expression values for barcodes are calculated (log2 of the ratio of RNA barcode cpm to DNA barcode cpm) in each replicate and collapsed into a mean per sample. Single barcodes with expression values ≥ 2 standard deviations apart from other barcodes for a given sequence in a given sample are dropped only from that sample, as this suggests the barcode exerted a cryptic regulatory effect (e.g. as a 3’UTR element) in the sample, or that either mutations in other barcode(s) on delivered plasmid or during sequencing preparation are contributing to spurious counts for that barcode. Penultimately, all barcodes for a sequence are dropped from an indivudal sample if the filtering steps up until now have resulted in 4 or fewer barcodes remaining for a given sequence in that sample.

After these filtering steps, expression is calculated on a per-barcode basis in each sample by taking log2 of the ratio of RNA cpm to DNA cpm. These values are then averaged for each sequence in each sample, resulting in a per-sequence expression value in each sequence. For the first assay, where the sole comparison is between alleles of a sequence, a student’s t-test is applied to compare the vector of samplewise mean expression values for one allele to those of the other allele. In the case of linear mixed modeling, as in the ATRA assay, the barcode-level expression values for the two respective alleles are used as model input, along with the corresponding allele for each barcode and drug/vehicle status of the corresponding samples to model both as expression ~ allele + drug + allele:drug. As a failsafe step, sequences with a calculable mean expression value in < 2 of the samples in a condition (of n=6 per condition in both experiments) were excluded from t-testing in the first experiment; in the ATRA experiment utilizing an LMM of barcode expression values, SNPs for which ≥ 60% of samplewise barcode expression measurements were missing (i.e., ≥ (12 replicates * 10 barcodes per allele * 2 alleles * 0.6 ==) 144 sample-barcode expression measurements) were excluded from analysis.

To perform statistical correction informed by the level of cross-sample and cross-barcode noise, the vector of Z values (t test) or F values (LMM) were stored for each SNP. For single-condition t-test analysis of the ATRA samples alone, Benjamini-Hochberg FDR correction was used, as our statistics of interest regarding these samples came almost entirely from the LMM (except for dual interaction and allele main effect cases, see below). To correct test statistics in the first assay, second assay vehicle-only, and LMM analyses, the same statistical comparison (t-test or LMM) as used for the sequences of interest was performed utilizing a vector of 10,000 (first assay), 20,000 (second assay LMM), or 5,000 (second assay vehicle samples alone) Z or F test statistics random “allelic” comparisons between elements that should have a known null difference in transcription—that is, 5,000-10,000 comparisons between two sets of 6 randomly selected barcodes—representing the median number of barcodes analyzed for a given sequence—from among the 110 barcodes assigned to the hsp68 promoter without human genomic sequences added upstream. This process effectively models the level of noise in expression calculations for the experiment due to e.g., variation in the cultures, across barcodes, and via biases during sequencing preparation. This method derives from an analogous statistical correction approach using non-targeting CRISPR gRNA sets from within a CRISPR screening library^37^.

For SNPs with significant allele and interaction coefficients in the LMM, a meaningful allele main effect was considered present if the single-condition vehicle and ATRA analyses showed the same allelic direction of effect, with a vehicle *q* < 0.1 and ATRA FDR < 0.1 (*i.e.,* near-significant within each condition of n=6, thus reasonably capable of achieving significance in the LMM analysis of the two conditions combined).

**MotifbreakR analysis**

In order to assess transcription factor binding motif perturbations corresponding to SNPs, we utilized the R package motifbreakR^38^ and its built in database of motif position-weight matrices (PWMs) from multiple public repositories. The tool by design identifies PWM matches overlapping input SNPs from dbSNP (here, version 151) for which at least one of the SNP alleles results in a genomic sequence significantly matching a given motif sequence. We used the default significance cutoff of p < 10^-4^ for calling motif matches in all analyses, and identified changes in motif match score using the tool’s default algorithm, which takes a weighted sum based on the position weights of each base in the motif sequence and considers these for the two alleles of the query SNP. As the use of dbSNP 151 required use of hg38, our queries thus took place in hg38 reference genome sequence space for the subset of SNPs with the same rsID in both the MPRA design and dbSNP 151. Computationally, all SNPs from a given set of positive and negative comparators were run through motifbreakR once with these settings to identify motif perturbations. For analysis of SNPs with identified in the first assay in N2a cells, SNPs with an allelic effect q-value < 0.05 were compared to those with an allelic effect q-value > 0.05; from the second assay, allele-ATRA interaction SNPs with q_interaction_<0.05 were compared to allele-main-effect-only SNPs (q_allele_<0.1 and q_interaction_>0.1), and allele main effect SNPs at q_allele_<0.1 were compared to the SNPs that were subject to neither main effects of allele or drug, nor their interaction (all q>0.1).

Frequencies at the level of TF (which can include several motifs) were considered as the number of SNPs matched to a given TF, regardless of the number or identity of motifs that SNP matched to. Null distributions of frequency were determined by 10,000 random selections of motif perturbations corresponding to *n* SNPs in the negative comparator SNP set, where *n* was the number of positive set SNPs analyzed based on the presence of an rsid in dbSNP 151. The p-value of frequency was then calculated from the empirical percentile of the positive SNP frequency count vs the distribution of frequencies in the negative set selections. Concordance was determined in a similar manner by permuting 10,000 random subsets of data covering *n* SNPs, but drawing motif assignments from the full set of SNPs analyzed and assigning a random “MPRA” allelic-directional effect to each SNP to compare observed rates of concordant allelic effects on motif match and reporter expression to the rates obtained by chance.

**CODE AVAILABILITY**

All code described in the analysis sections, including an MPRA analysis function and the described bash scripts for extracting barcode sequences from raw fastq sequencing files, are available at <https://bitbucket.org/jdlabteam/n2a_atra_mdd_mpra_paper/src/master/>

**SUPPLEMENTAL FIGURES AND TABLES**

**Supplemental Figure 1.** A) Cross-replicate comparison of individual barcode counts per million (CPM). B) Cross-replicate comparison of element-level expression (mean barcode RNA/DNA ratio). Pearson correlation coefficients are shown above the diagonal in both panels.

**Supplemental Figure 2.** Extended motifbreakR results from the first MPRA, showing enrichment analysis results agnostic to the motifbreakR-defined “strength” of motif change between alleles (left-hand columns). A) TFs with enriched frequency of motifs among functional SNPs. The corresponding number of functional SNPs matched to each TF for a given strength are shown. B) TFs with greater than predicted concordant motif and MPRA effects among functional SNPs. Concordant effects were defined by greater MPRA expression driven by the allele better-matched to the corresponding TF motif and vice versa—the expected behavior of strictly activating TFs.

**Supplemental Figure 3.** Additional functional SNPs corresponding to an enriched TF motif group (NR3C1) from the first assay. Replicates are indicated by individual points and their connecting lines.

**Supplemental Figure 4.** Extended motifbreakR results from the ATRA treatment MPRA, showing enrichment analysis results agnostic to the motifbreakR-defined “strength” of motif change between alleles (left-hand columns). A) TFs with enriched frequency of motifs among retinoid-independent functional SNPs compared to SNPs with no detected allelic effects. The corresponding number of functional SNPs matched to each TF for a given strength are shown. Heatmap is split into two vertical slices for visibility. B) TFs with motifs overrepresented among ATRA-dependent (interaction) functional SNPs compared to ATRA-independent functional SNPs. The heatmap is shown in halves for visibility. TFs identified as ATRA-upregulated in human neuroblastoma lines^39^ are in bold font.

**Supplemental Table 1.** SNPs with functional effects detected in at least one analysis, GRCh19 and GRCh38 coordinates, corresponding GWAS tag variant, and their effect calculated as alt vs. ref allele in the first assay and second assay single-conditions.
