## Supplementary figures and images for "Transcriptional-Regulatory Convergence Across Functional MDD Risk Variants Identified by Massively Parallel Reporter Assays"

### Figure S1

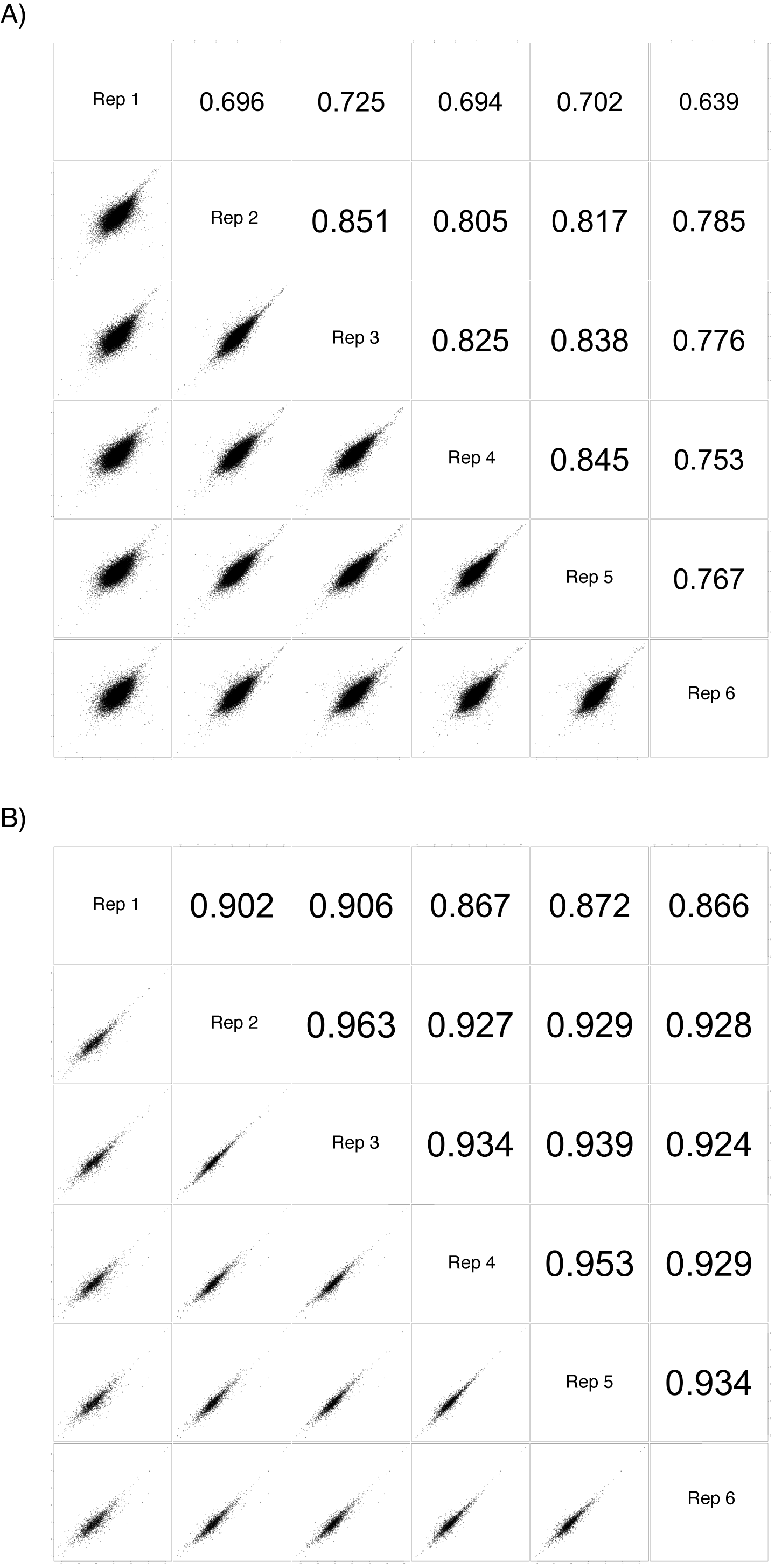

### Figure S2

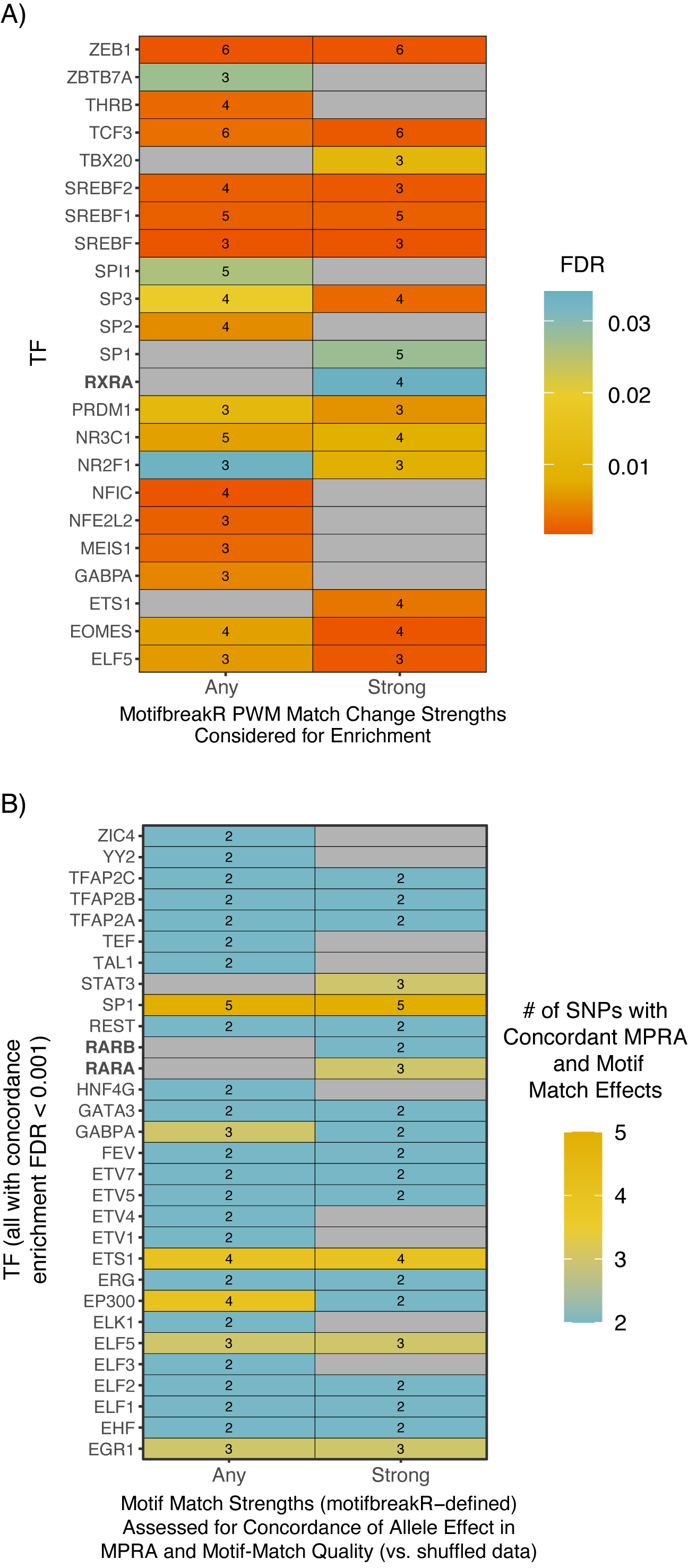

### Figure S3

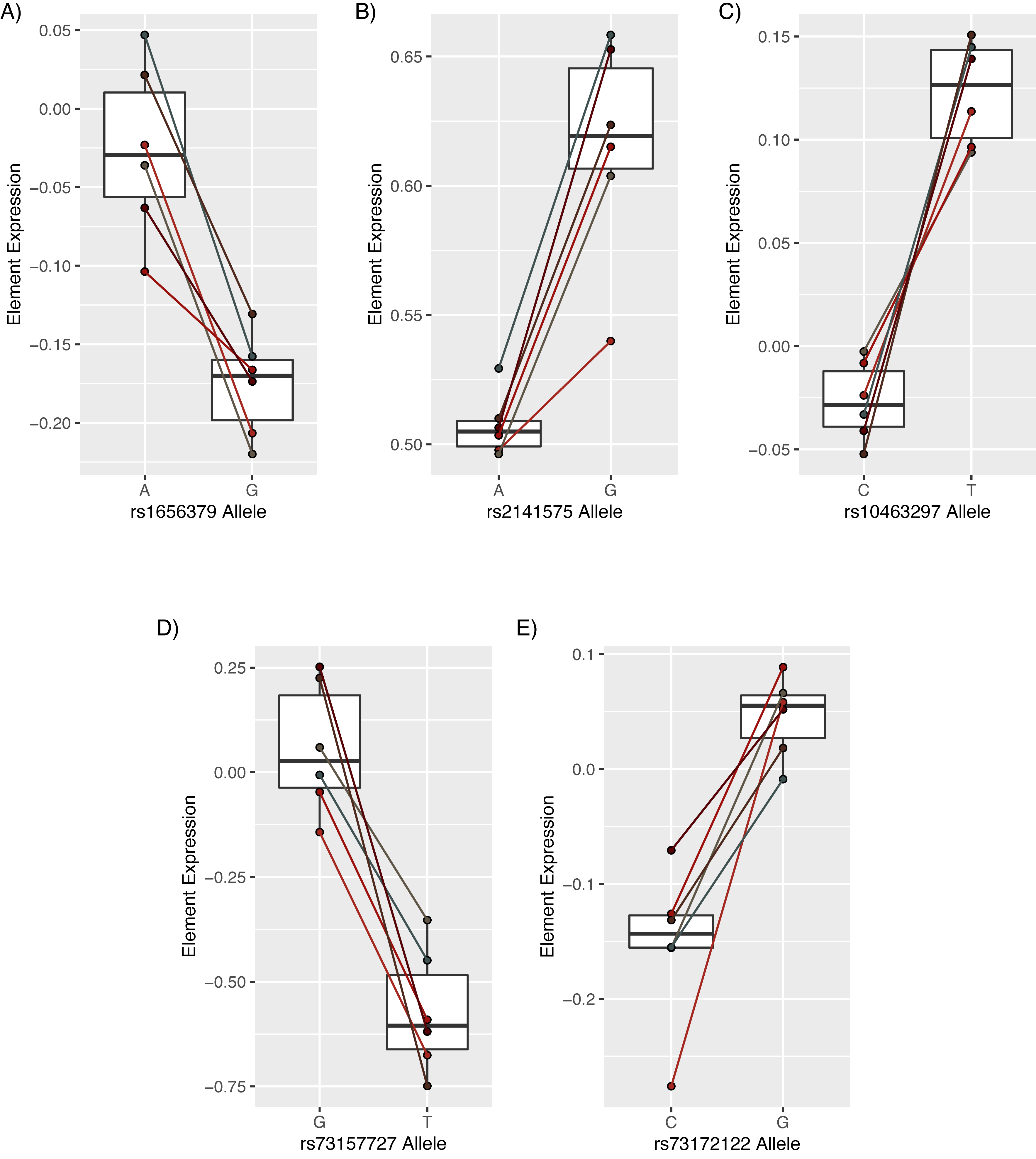

### Figure S4

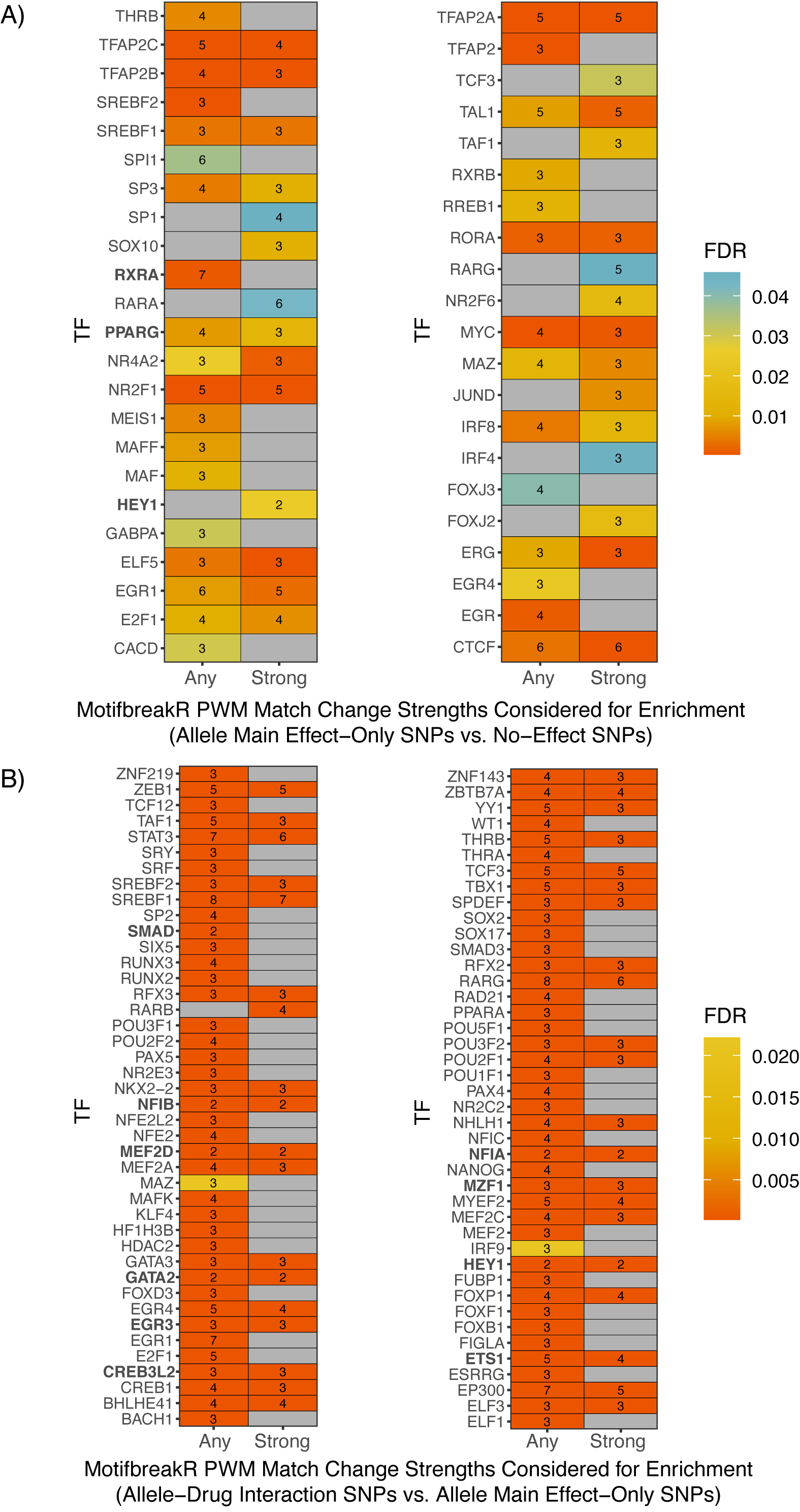
